# Thalamocortical Spindles occur during Rapid Eye Movement Sleep in Mouse Somatosensory Pathway

**DOI:** 10.1101/2023.10.26.564196

**Authors:** Flore Boscher, Katlyn Jumel, Tereza Dvořáková, Luc Gentet, Nadia Urbain

## Abstract

Rapid eye movement sleep (REM) is often considered as a homogeneous state of sleep. However, the frequent occurrence of transient events indicates that it may be separated into two distinct, phasic and tonic, substates. During tonic REM, we found local appearances of spindle waves in the barrel cortex concomitant with strong delta power on the local field potential. Subthreshold spindle oscillations in neurons of the ventral posterior medial nucleus further confirmed the thalamic origin of these cortical spindles. Spindle oscillations were suppressed in phasic REM, while thalamus spike firing increased associated with rapid whisker movements of mice and cortical activity transitioned to an activated state. During REM, sensory thalamus and barrel cortex therefore alternate between high (wake-like) and low (non-REM sleep-like) activation states, possibly allowing transient sensory integration windows to emerge throughout this paradoxical sleep stage.

## INTRODUCTION

Mammalian sleep includes two main states: slow wave sleep, or non-rapid eye movement sleep (NREM), and the subsequent REM sleep (REM). REM has long been defined as a state of fast, desynchronized and low voltage electroencephalogram (EEG) resembling wakefulness, with the occurrence of transient bursts of ocular movements coupled with a complete muscle atonia ^1,2^. This striking EEG activation during a sleep state inspired Jouvet to coin REM with the name “paradoxical sleep” ^3^. Many further transient events co-occurring with rapid eye movements, including muscle twitches, breathing fluctuations, heart rate surges or facial movements, associated with electrophysiological markers such as pontine P-waves or an increase in EEG and hippocampal theta frequency, have since been described ^4–8^. These phasic events appear on the background of an apparently more quiescent, tonic state, suggesting that REM is not an homogeneous sleep state ^9–11^. Indeed, local field potential (LFP) recordings of brain areas revealed the unexpected presence of local delta oscillations during REM both in humans and rodents ^12–15^. Delta activity had mostly been described in NREM sleep, and more recently in wakefulness ^16–19^. Interestingly, NREM delta waves, along with spindles and ripples, have been shown to be involved in mnemonic processes ^20,21^. Since memory consolidation and emotional processing also have been functionally ascribed to REM sleep ^22–28^, it raises the question whether delta oscillations play a similar mnemonic function in REM. The activation of a thalamocortical network including limbic and parahippocampal areas was reported during rapid eye movements in humans ^29–33^ but still very little is known about the neuronal mechanisms subtending these observations. Considering that, except in early stages of the development, spindle-like waves in REM were rarely reported in the literature ^34,35^, it was proposed that hippocampal Theta rhythm in REM may assume this role in mnemonic processes ^27,36^.

The present study aimed at exploring cortical oscillations and their thalamic correlates associated with REM substates. Focusing on the mouse somatosensory system, we report during REM a highly dynamic interplay between local oscillations in the primary somatosensory (S1) cortex and membrane potential dynamics and neuronal firing activity of the thalamic ventral posterior medial nucleus (VPM). We demonstrated 1- that S1-LFP displayed features of NREM sleep during tonic REM, characterized by strong delta (2 - 6 Hz) power associated with spindles, which are suppressed during phasic periods of whisker movements, when S1-LFP transitioned to an activated state, and 2- its thalamic neuronal correlates which confirm the thalamic origin of the observed REM-spindles.

## RESULTS

We performed S1-LFP associated with extra- and intracellular recordings of thalamic relay cells while simultaneously measuring the EEG and the electromyogram (EMG). These recordings allowed us to investigate the cortical dynamics and their thalamic correlates in REM and to compare them with those observed in wakefulness (WK) and NREM sleep in non-anesthetized head-fixed mice.

### Rapid whisker movements are associated with phasic cortical activity in REM

Rapid whisker movements (RWM) were observed during REM, as we previously reported ^37^ (Figure 1). RWM represented on average 20%, while non-whisking periods (REM-noWh) accounted for about half of the total REM duration (Figure 1B, average across 110 episodes of REM from 32 mice); and the probability to observe RWM was significantly higher in the last two thirds of REM episodes compared with the first third (respectively 4.3 ± 0.2 %, 6.4 ± 0.3 %, and 6.6 ± 0.4 % in the first, second, and third parts of the REM). Compared to free-whisking during wakefulness (Wh), RWM were faster (RWM, 14.0 ± 0.3 Hz vs Wh, 10.6 ± 0.2 Hz; n= 35 mice) and smaller in amplitude (RWM, 34.1 ± 0.6° vs Wh, 39.2 ± 0.8°) (Figure 1C-E). Whisking prominence, i.e. the amplitude of protraction/retraction cycles, was significantly lower in REM (RWM, 14.6 ± 0.2° vs Wh, 21.1 ± 0.6°), indicating a more focal whisking than in wakefulness. RWM episodes were also shorter than in wakefulness (RWM, 1.05 ± 0.04s vs Wh, 1.53 ± 0.07s), but with no variation in the number of cycles per episode (RWM, 12.3 ± 0.5 vs Wh, 14.2 ± 0.7).

**Figure 1.**
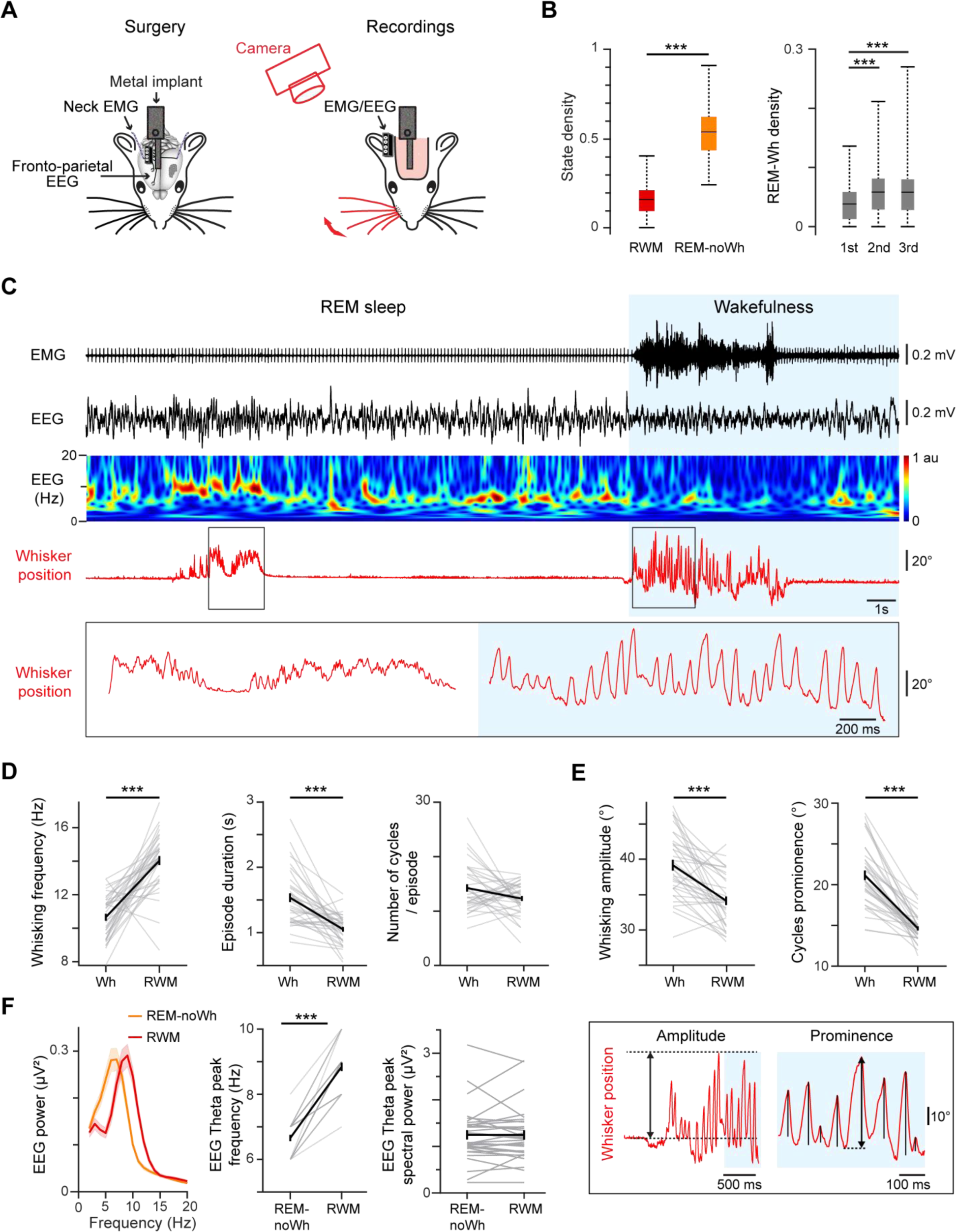
Phasic activity in REM sleep. (A) Schematic of the experimental set-up. Left, EEG and EMG electrodes were implanted and a metal head-holder was stereotaxically glued to the skull. Right, whisker movements were filmed with a high-speed camera in parallel of EEG and EMG monitoring. (B) Left, percentage of time spent in RWM and REM-noWh during REM episodes (see Methods for calculations). Right, percentage of time spent in RWM during the first, second and third part of REM episodes longer than 60s (ANOVA for repeated measures). Boxplots indicate mean (black line) and interquartile range. (C) Example of a typical EEG recording during REM and the subsequent awakening. Top to bottom: neck EMG, EEG, corresponding EEG wavelet time-frequency spectrum and whisker position (in red) with the enlarged view of the black boxes. Whisker movements are indeed not restricted to wakefulness. (D) From left to right: whisking frequency, duration and number of protraction/retraction cycles per episode, in wakefulness and REM. (E) Average whisking amplitude (left) and prominence (right) per mouse, in wakefulness and REM, with an illustration of the method used to estimate these parameters (bottom). (F) Grand average power spectral density (PSD) of the EEG during REM sleep with (red) or without (orange) whisker movements (Left), and corresponding averaged EEG theta peak frequency (Middle) and spectral power (right) per mouse. B, F, n= 32 mice; data were compared using Wilcoxon or Friedman tests. D-E, n= 35 mice; *t*-test for paired data. Mean ± sem in black. Note that y axes do not start at “0”.

RWM were associated with an increase in EEG theta frequency compared to tonic REM (RWM, 8.8 ± 0.1 Hz vs REM-noWh, 6.7 ± 0.1 Hz; average across 32 mice) but with no change in theta power (Figure 1F). To examine the hippocampal activity associated with RWM, we performed LFP recordings in the CA1 area of the dorsal hippocampus (HPC) (Figure S1); in all animals (3 mice), we observed that hippocampal theta oscillations show similar properties and a high correlation with the ones on the EEG. These results support the hypothesis that EEG theta rhythms mostly reflect hippocampal REM activity in mice ^38,39^. RWM were also associated with an increase in gamma activity at both the EEG and hippocampal level: gamma oscillations (60 - 90 Hz) power was increased during RWM, and gamma amplitude was modulated on the theta wave (Figure S1D-F). Taken together, our results indicate that RWM are strongly correlated with phasic hippocampal activity during REM. We will therefore hereafter consider two substates of REM: RWM and REM-noWh, corresponding to previously-reported phasic and tonic REM substates ^9–11^.

### S1-LFP exhibits both wakefulness and NREM features in REM

Quiet wake periods (qWK) characterized by large-amplitude slow fluctuations in the delta range (2-6 Hz) in S1-LFP, alternated with active wake episodes, either with (Wh) or without (aWK) whisking, as we described previously ^18^ (Figure 2A-C; delta power: qWK, 4.6 ± 0.7 µV²; Wh, 0.8 ± 0.1 µV²; aWK, 0.9 ± 0.2 µV²). Similarly, a high delta power was observed in S1-LFP during tonic REM (4.5 ± 0.7 µV²), which decreased significantly during RWM bouts (2.8 ± 0.5 µV²; n= 28 mice). Interestingly, strong delta power in tonic REM was restricted to S1-LFP (S1-LFP: 3.3 ± 0.8 µV², M1-LFP: 1.3 ± 0.3 µV², EEG: 1.3 ± 0.2 µV²; n= 11 mice) in contrast with the more global delta rhythm observed in qWK (S1-LFP, 4.0 ± 0.7 µV²; M1-LFP, 4.6 ± 0.8 µV²; EEG, 2.3 ± 0.2 µV) (Figure S2A-D). These results suggest that S1-LFP displays, in REM, delta activity that closely resembles the one observed during wakefulness, with a decrease in delta power associated with whisking in both vigilance states.

**Figure 2.**
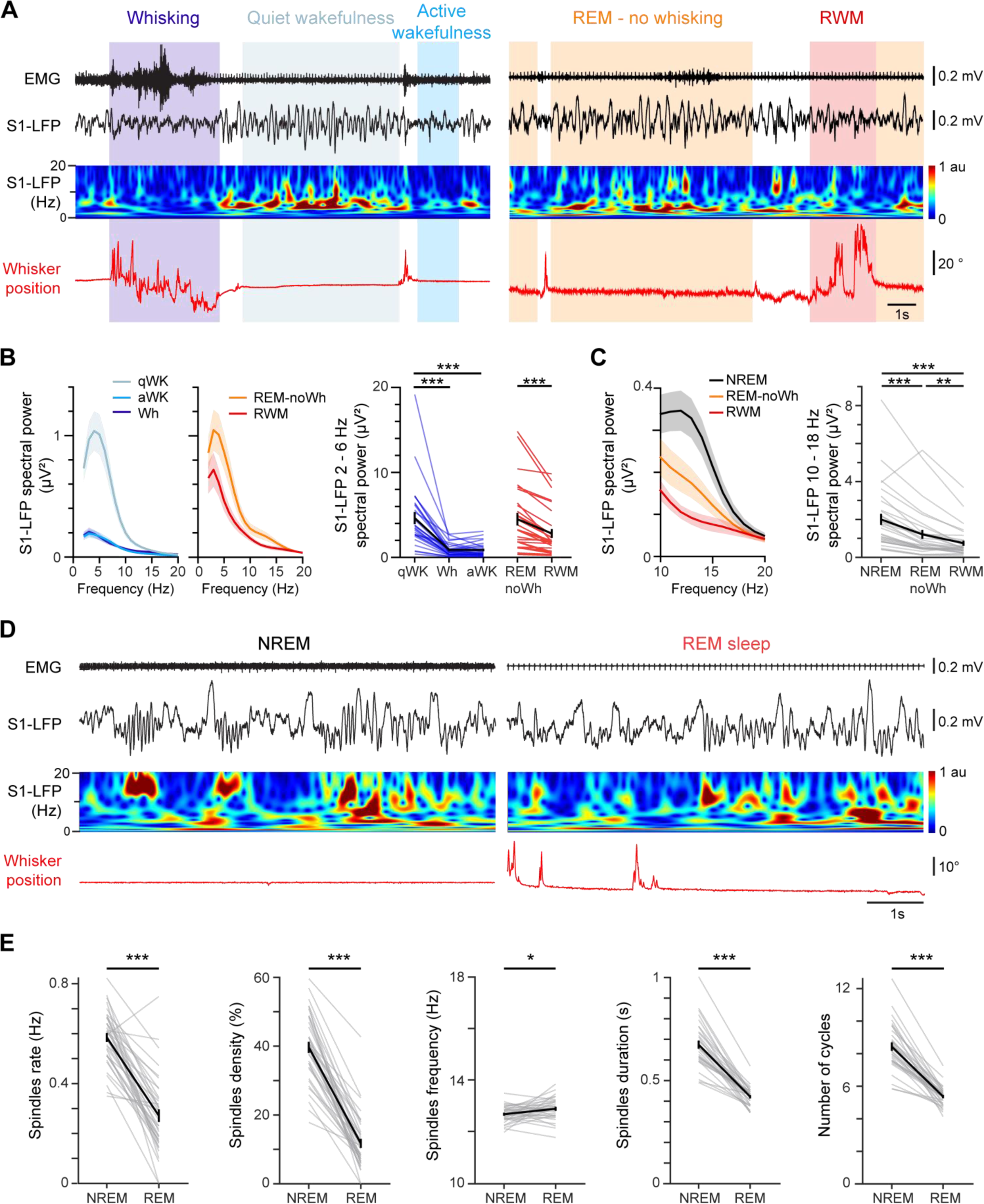
S1-LFP exhibited in REM both typical wakefulness and NREM features. (A) S1-LFP activity during wakefulness (left) and REM (right). Top to bottom: neck EMG, S1-LFP, corresponding S1-LFP wavelet time-frequency FFT (color graph) and whisker position (in red). (B) Left: grand average PSDs of S1-LFP in the different substates of wakefulness (blue) and REM (red). Right: corresponding average S1-LFP spectral power in the delta range (2 - 6 Hz; n= 28 mice). (C) Left: grand average PSD of S1-LFP in the sigma range (10 - 18 Hz) in NREM (black), REM-noWh (orange) and RWM (red) (n= 28 mice). Right: corresponding average S1-LFP spectral power per mouse. (E) S1-LFP activity during NREM and REM. Top to bottom: same as (A). (E) Average spindles parameters per mouse in NREM and REM. From left to right: rate and density (n= 37 mice), frequency, duration and number of cycles (n= 32 mice). The solid line represents the mean ± SEM; data were compared across states using Friedman and Wilcoxon tests (C-D), and *t*-test for paired data (E).

Strikingly, S1-LFP also displayed a strong sigma (10 - 18 Hz) power during REM, well known as a hallmark of NREM spindle activity (Figure 2C; REM, 1.3 ± 0.3 µV² vs NREM, 2.2 ± 0.3 µV²; n= 28 mice). In order to determine whether this REM-sigma activity was subtended by spindles, we applied an algorithm that was developed during a previous study of S1-LFP in NREM ^40^ to automatically detect individual spindles (Figure S2E; see Materials and Methods). The algorithm robustly detected spindles throughout REM (Figure 2D-E; S2E-H), but with a lower rate and density than those observed in NREM (Rate: REM, 0.27 ± 0.03 vs NREM, 0.58 ± 0.02 spindles/s; density: REM, 11.7 ± 1.3 vs NREM, 39.6 ± 1.6 %, n= 37 mice). Spindles were of similar frequency in REM than in NREM (REM, 12.9 ± 0.1 Hz vs NREM, 12.7 ± 0.1 Hz; n= 32 mice), but of shorter durations (REM, 0.42 ± 0.01 s vs NREM, 0.67 ± 0.02 s), with a lower number of cycles per spindle (REM, 5.4 ± 0.1 vs NREM, 8.4 ± 0.3). They occurred preferentially outside RWM.

Since delta waves are also typically observed in NREM ^40,41^, we calculated the delta band power on S1-LFP in REM-noWh compared with NREM (Figure S2H; REM-noWh, 4.5 ± 0.7 µV² vs NREM, 5.2 ± 0.8 µV²; n= 28 mice). On normalized spectra of S1-LFP between 2 and 20 Hz, REM-noWh delta power was significantly larger than in NREM (REM, 56.1 ± 1.3% vs NREM, 50.0 ± 1.6%).

Taken together, these data indicate that REM is composed of two distinct substates with characteristics reminiscent of both wakefulness and NREM: a phasic state characterized by a cortical activation associated with RWM, similar to wakefulness; and tonic REM, characterized by delta oscillations coupled with the appearance of spindles, closely resembling S1-NREM.

### Thalamic neurons display both wakefulness and NREM activity patterns during REM

In order to further probe the REM substate dynamics observed on S1-LFP, we recorded the firing activity and membrane potential (Vm) of somatosensory relay cells in the ventro-posterior medial (VPM) thalamus. VPM neurons mean firing rate increased significantly during active compared to quiet wakefulness, and further increased during whisking (Figure 3A-B; Wh, 20.3 ± 2.3 spikes/s, aWK, 12.1 ± 2.0 spikes/s and qWK, 5.6 ± 1.4 spikes/s; n= 32 cells), in agreement with our previous study ^18^. Those same neurons exhibited a similar increase in mean firing rate associated with RWM during REM (24.8 ± 3.4 vs REM-noWh, 16.2 ± 2.0 spikes/s). VPM cells were as active during RWM as in wake whisking (Figure 3B, n= 35 cells). Moreover, around half of the cells recorded in both wakefulness and REM (17/32) significantly increased their firing rate during whisking in wakefulness, and 70% of them were similarly modulated by RWM (Figure 3C). Less than 10% were modulated in RWM only. Self-generated whisker movements may therefore drive thalamic activity through direct sensory activation in REM, as previously described in wakefulness ^18,42^.

**Figure 3.**
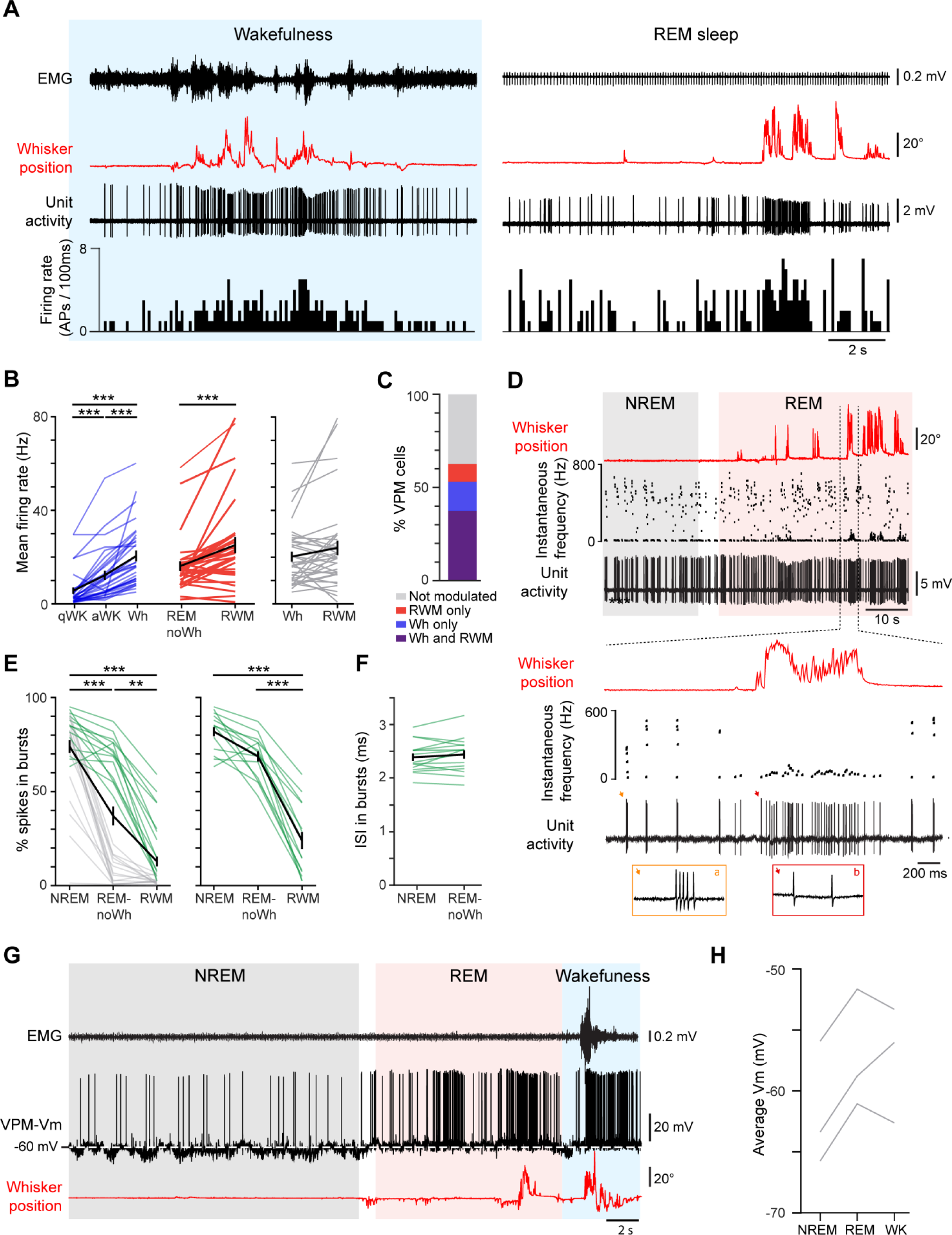
Thalamic neurons activity in REM exhibited both wakefulness and NREM patterns. (A) Activity of a typical VPM cell during wakefulness (left) and REM (right). Top to bottom: neck EMG, whisker position (in red), VPM neuron unit activity and firing rate (100ms bins). (B) Left: mean firing rate of VPM neurons in wakefulness (blue) and REM (red; n= 32 cells recorded in all five states). Right: comparison between whisking mean firing rates in wakefulness and REM (n= 35 cells in Wh and RWM). (C) Percentage of whisking-modulated VPM cells (n= 32 cells). (D) Top: unit activity of an example VPM neuron showing bursting activity across NREM and REM, and associated whisker position (red trace). Middle: the instantaneous frequency (1/interspike interval (ISI)) showing that high-frequency bursts were observed in both NREM and REM, but outside of whisker movements. Bottom: enlarged view of the boxed area with 20ms insets showing a burst of spikes in REM-noWh or a single-spike firing in RWM. (E) Percentage of spikes in bursts in NREM, REM-noWh and RWM for all cells (left, n= 29 recorded in all three states) and for the subpopulation of VPM neurons that fired bursts of spikes in REM-noWh (green; right; n= 13). (F) Mean duration of ISIs within bursts (n= 16 cells exhibiting bursts in NREM and REM-noWh). (G) Membrane potential (Vm) of a typical VPM cell and associated whisker position (in red) across NREM, REM and wakefulness. (H) Average Vm per cell in NREM, subsequent REM and wakefulness. B, F, Black line, mean ± SEM; data were compared across states using ANOVA for repeated measures or *t*-test for paired data.

While all recorded VPM cells fired, as expected, high frequency bursts of spikes during NREM, 43% (16/37) of them conserved this firing pattern during REM, and only switched to a tonic firing mode during RWM (Figure 3D-E; NREM, 82.1 ± 3.0 % of spikes included in bursts; REM-noWh, 70.3 ± 2.8 %; RWM, 25.4 ± 5.3 %; n= 13 cells). Bursts of spikes in REM displayed the same characteristics as the NREM thalamic bursts ^40,43^, i.e., same mean interspike interval (ISI; Figure 3F; 2.4 ± 0.0 ms in both NREM and REM; n= 16 cells), and same dynamics of ISI durations and spike amplitudes within a burst (Figure S3A). They occurred at the same rate than during NREM (Figure S3B; REM, 1.2 ± 0.5 vs NREM, 1.1 ± 0.4 burst/s), despite an increase in burst duration (REM, 8.0 ± 0.1 ms vs NREM, 6.0 ± 0.1 ms), with a higher number of spikes per burst (REM, 4.3 ± 0.0 vs NREM, 3.5 ± 0.0 spikes). Finally, membrane potential recordings showed that VPM-Vm was more depolarized during REM than NREM (Figure 3G-H; 4.5 mV more depolarized on average; n= 3 cells for which our intracellular recordings were stable across the entire sleep-wake cycle), in agreement with a previous study in cats ^44^. Together, our data show a highly dynamic modulation of thalamic cells firing during REM: a tonic activity during RWM, reminiscent of whisking in wakefulness, as well as high frequency bursts, which closely resemble their NREM firing pattern.

### Thalamocortical coupling is maintained during REM in the somatosensory pathway

High frequency VPM thalamic bursts occur preferentially on the rising phase of cortical spindle cycles, and were therefore believed to be directly involved in spreading spindles generated within the thalamus to the cortex during NREM ^40,45^. To obtain further insight into the thalamic and cortical coupling occurring during REM, we analyzed the timing of VPM spikes relative to S1-spindle cycles in in REM. We found that fewer VPM spikes were phase-locked in REM compared to NREM (17/24 versus 22/24, respectively; Rayleigh test p< 0.01; Figure 4A-C). However when the neurons were phase locked, they showed similar phase preference (NREM, 83 ± 14° before the spindle trough; REM, 81 ± 11°; n= 17 cells). Reciprocally, VPM population spike-triggered averaging (STA) on S1-LFP revealed a peak, with a lag of less than 20ms after thalamic spiking in both NREM and REM (Figure 4D; REM, 19ms vs NREM, 15ms). Remarkably, our intracellular recordings revealed the presence of spindle oscillations on the Vm of a subset of VPM neurons (3/6) in NREM as well as in REM (Figure 4E). Thalamic Vm spindles density decreased in REM compared to NREM, concomitantly with the observed decrease at the level of S1-LFP (NREM, 35% vs REM, 5%; n= 3 cells).

**Figure 4.**
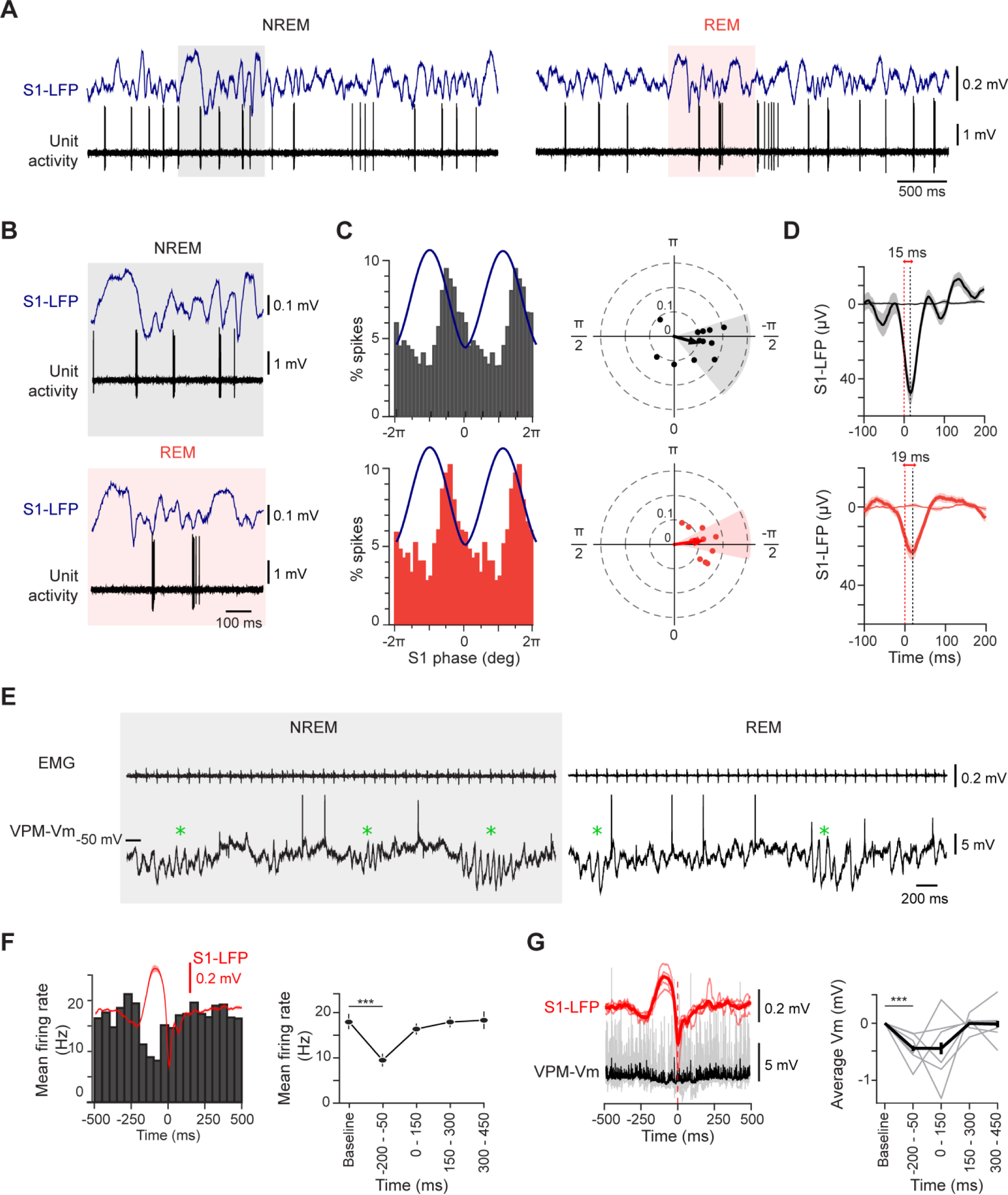
Thalamo-cortical dynamics in NREM and REM. (A) VPM unit activity and S1-LFP during spindle oscillations in NREM (left) and REM (right). (B) Enlarged view of the colored areas in (A) corresponding to a spindle oscillation in S1-LFP in NREM (top) and REM (bottom). (C) Left, grand average spike distribution of VPM cells on S1-LFP spindle cycles (in blue; 18 bins/cycle) in NREM (top) and REM (bottom). Right, corresponding spike firing preferred phases and modulation amplitudes on spindle cycles for modulated VPM cells. Only cells that were modulated during S1-LFP spindle cycles in both NREM and REM were considered (p < 0.01 Raleigh test, n= 17 cells). (D) S1-LFP grand average (mean ± SEM) triggered on VPM spikes occurring during NREM (top) or REM (bottom; n= 20 cells; thin lines, surrogates). (E) Example of Vm of a thalamic cell recorded in NREM (left) and REM (right) together with EMG. Oscillations in the spindle frequency range (green stars) can be observed in both states. Spikes are truncated. (F) Spindles preceded by a delta wave occurring in REM were detected on S1-LFP and aligned on the spindle onset. Extracellular spike time occurrences were collected for VPM cells (n= 22), Peristimulus time histograms (PSTHs) were computed and a grand average was calculated (left; 50ms bins; n= 22 cells); the corresponding grand average S1-LFP delta oscillations is superimposed in red and the grand average firing rates (mean ± sem) is illustrated to the right. (G) Left, S1-LFP and corresponding normalized Vm population averages for VPM cells (n= 6) aligned on cortical spindles onset. Right, average normalized median Vm (150 ms bins) for these VPM cells (Black, mean ± sem). The baseline Vm (−500 to −300 ms before S1-spindles onset) was subtracted to get the normalized data). Data from each bin were compared to baseline using a *t*-test for paired data (F) or a Wilcoxon test (G). A-D, NREM in black, REM in red.

In addition, we observed that spindles in S1 were often preceded by a delta wave during REM, as previously reported during NREM ^40^. Since these delta events in NREM were associated with a decrease in VPM firing rates, we examined the firing rate of VPM cells recorded extracellularly relative to S1-spindles preceded by delta waves during REM. We computed peristimulus time histograms (PSTHs) of VPM spikes aligned to spindle oscillation onset and found a significant decrease in VPM discharge during delta events (Figure 4F; −200 to −50 ms, 11.4 ± 1.7 Hz vs baseline, 20.6 ± 1.7 Hz; n= 22 cells). At the subthreshold level, these REM delta events were accompanied by a hyperpolarization of VPM-Vm (Figure 4G; −200 to −50ms period: −0.4 ± 0.1 mV; n= 6 cells). Taken together, these results suggest a strong thalamocortical oscillatory coupling during REM, with marked similarities to NREM coupling.

**Figure S1.**
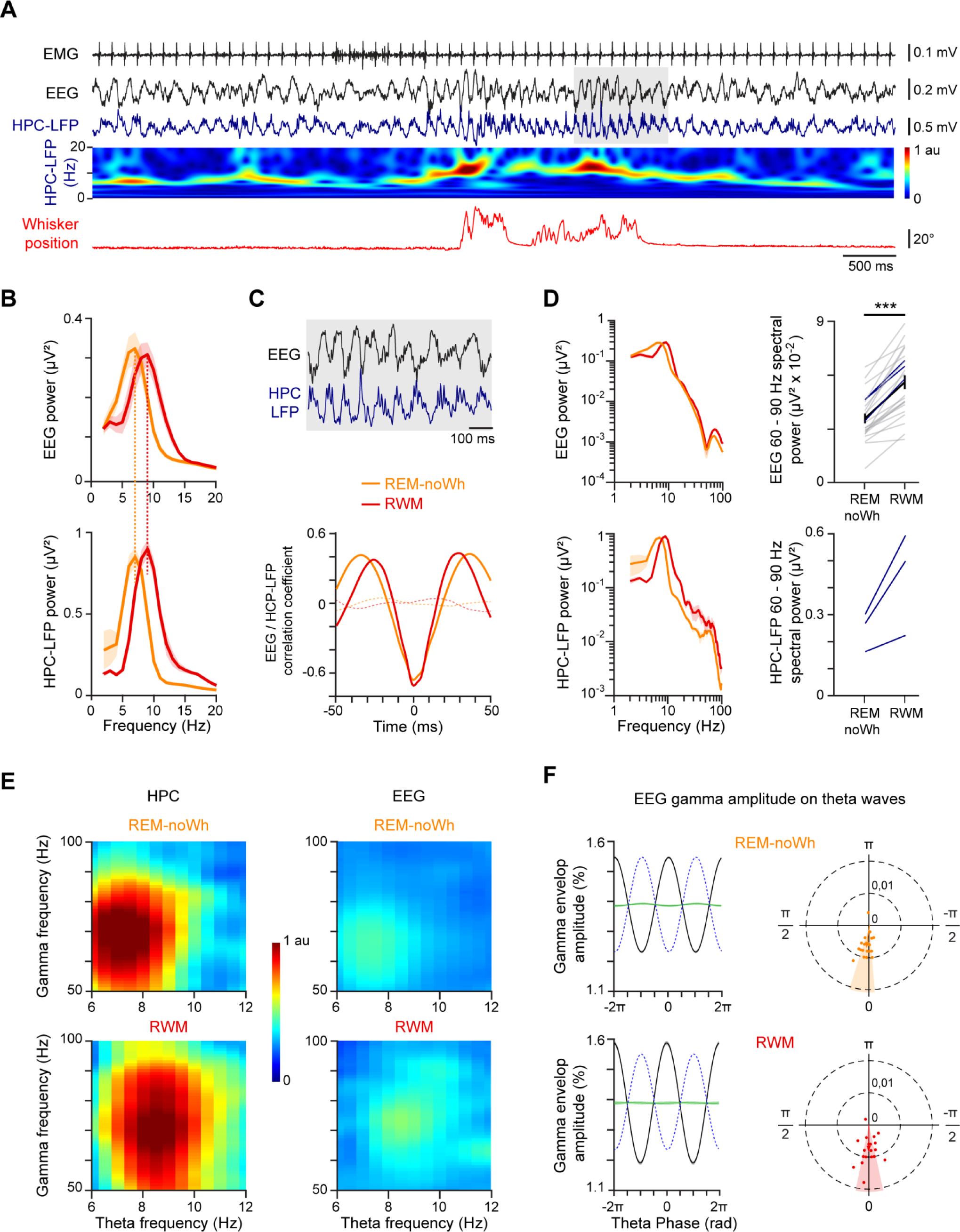
Hippocampal activity is highly correlated to the EEG in the theta band in REM. (A) Typical recording of HPC activity during and outside whisking behavior in REM. Top to bottom: neck EMG, EEG, HPC-LFP (blue) and corresponding HPC-LFP wavelet time-frequency spectrum (color graph) and whisker position (red). (B) Grand average PSDs of EEG (top) and HPC-LFP (bottom) recorded on 3 mice across REM-noWh (orange) and RWM (red). (C) Top, enlarged view of the grey box in (A). Bottom, grand average cross-correlogram between EEG and HPC-LFP in REM-noWh and RWM, compared to shuffled data (dashed lines). (D) Left, grand average PSD of EEG (top) and HPC-LFP (bottom) in REM-noWh and RWM displayed with a logarithmic scale. Right, corresponding average EEG (mean ± sem in black; Wilcoxon test) and HPC-LFP spectral power per mouse in the gamma range (60 - 90 Hz); blue lines correspond to the 3 cells for which both EEG and HPC-LFP recordings were available. (E) Phase-amplitude comodulograms for HPC-LFP (left) and the EEG (right) in REM-noWh (top) and RWM (bottom; see Methods). Warm colors illustrate that the amplitude of gamma oscillations is modulated with the theta wave. (F) Left, distribution of the mean EEG gamma envelop amplitude (black line) on the EEG theta waves (dotted blue line) in REM-noWh (top) and RWM (bottom). Average of shuffled gamma envelops is represented in green. Right, gamma oscillation maximum amplitude preferred phases and modulation amplitudes on theta waves for the EEG, in REM-noWh and RWM. Colored areas represent the confidence interval at 95%. (B-F) REM-noWh data are illustrated in orange and RWM data in red. C-F, HPC, n= 3 mice; EEG, n= 23 mice.

**Figure S2.**
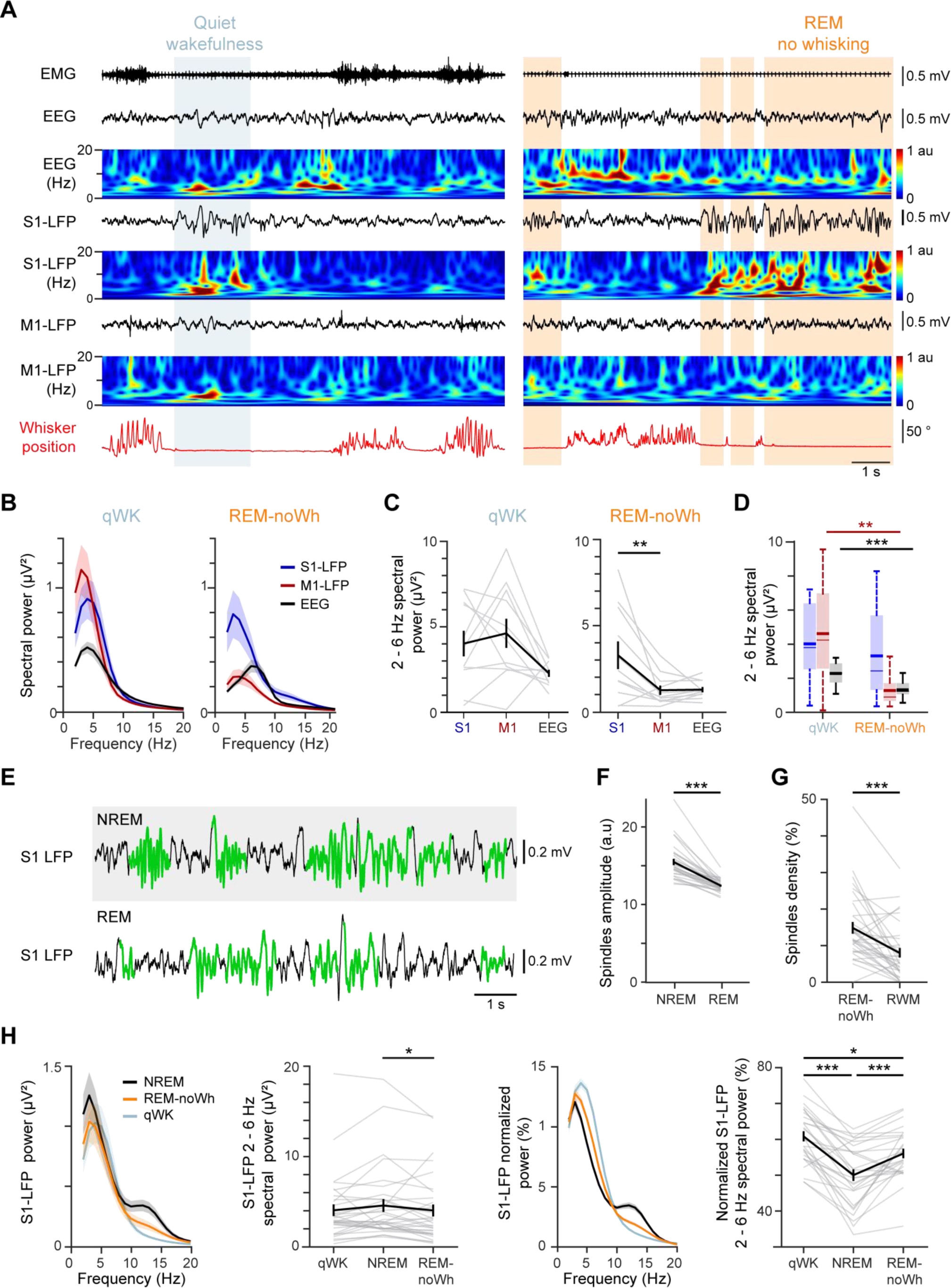
Delta and spindle oscillations in REM versus wakefulness and NREM. (A) Cortical activity during wakefulness and REM. Top to bottom: neck EMG, EEG, S1-LFP, M1-LFP and their corresponding wavelet time-frequency FFTs (color graphs) and associated whisker position (in red). (B) Grand average PSD of S1-LFP (blue), M1-LFP (red) and EEG (black) in qWK (left) and REM-noWh (right). (C) Average S1-LFP, M1-LFP and EEG spectral power per mouse in the delta range (2 - 6 Hz) in qWK (left) and REM-noWh (right; in black, mean ± sem). (D) Comparison of grand average S1-LFP (blue), M1-LFP (red) and EEG (black) spectral powers in the delta range between qWK and REM-noWh. Boxplots indicate mean (thick line), median (thin line) and interquartile range. (thick line, mean; thin line, median). B-D, n= 11 mice. (E) Example of spindles detection (in green) in S1-LFP by the algorithm, in NREM (top) and REM (bottom). (F) Average spindles relative amplitude in NREM and REM (see Methods, n= 32 mice). (G) Average spindles density calculated in REM-noWh and RWM (n=35 mice). (H) Left, Grand average PSD of S1-LFP in NREM (black), REM-noWh (orange) and qWK (grey) and corresponding spectral power in the delta range. Right, Distribution of grand average spectral power (normalized PSD) of S1-LFP in NREM, REM-noWh, and qWK and corresponding mean distribution per mouse in the delta range (in black, mean ± sem; n= 28 mice). C-H, data were compared using Friedman or Wilcoxon tests.

**Figure S3.**
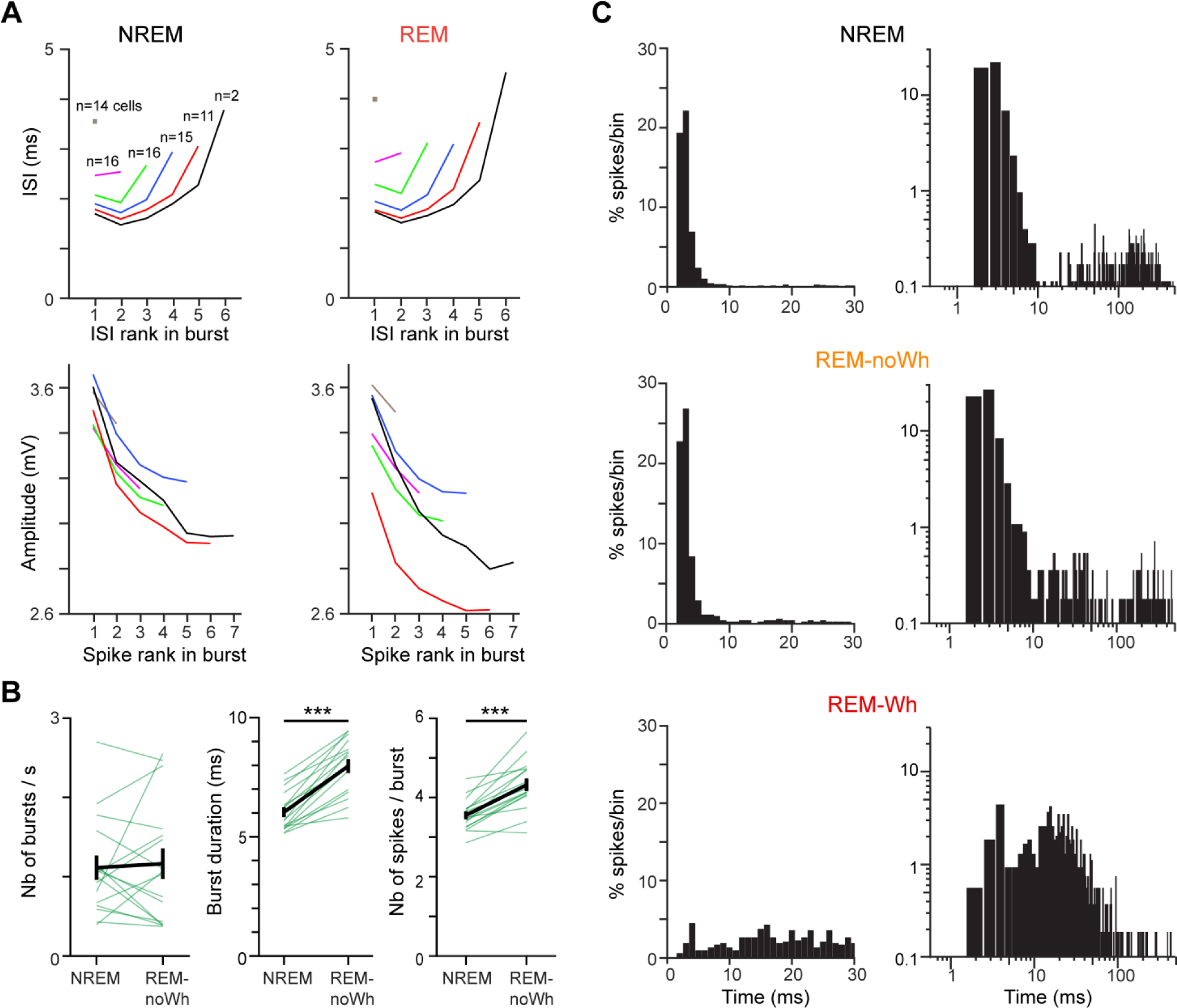
A subpopulation of VPM neurons exhibit a similar firing pattern in REM and NREM. (A) ISI (Top) and spike amplitudes (bottom) plotted as a function of the spike rank within a burst in NREM (left) and REM (right; n= 16 cells). (B) From left to right: average bursts rate, duration, and number of spikes, per cell (n= 16). (C) Example interspike interval histograms (ISIHs, 1 ms bins) in NREM (top), REM-noWh (middle) and RWM (bottom) for a VPM cell that fires bursts of spikes in REM-noWh; x and y axis are drawn in linear (left) and log10 (right) scales. The total number of spikes (100%) is the number of spikes collected to compute 0-500 ms ISIHs.

**Figure S4.**
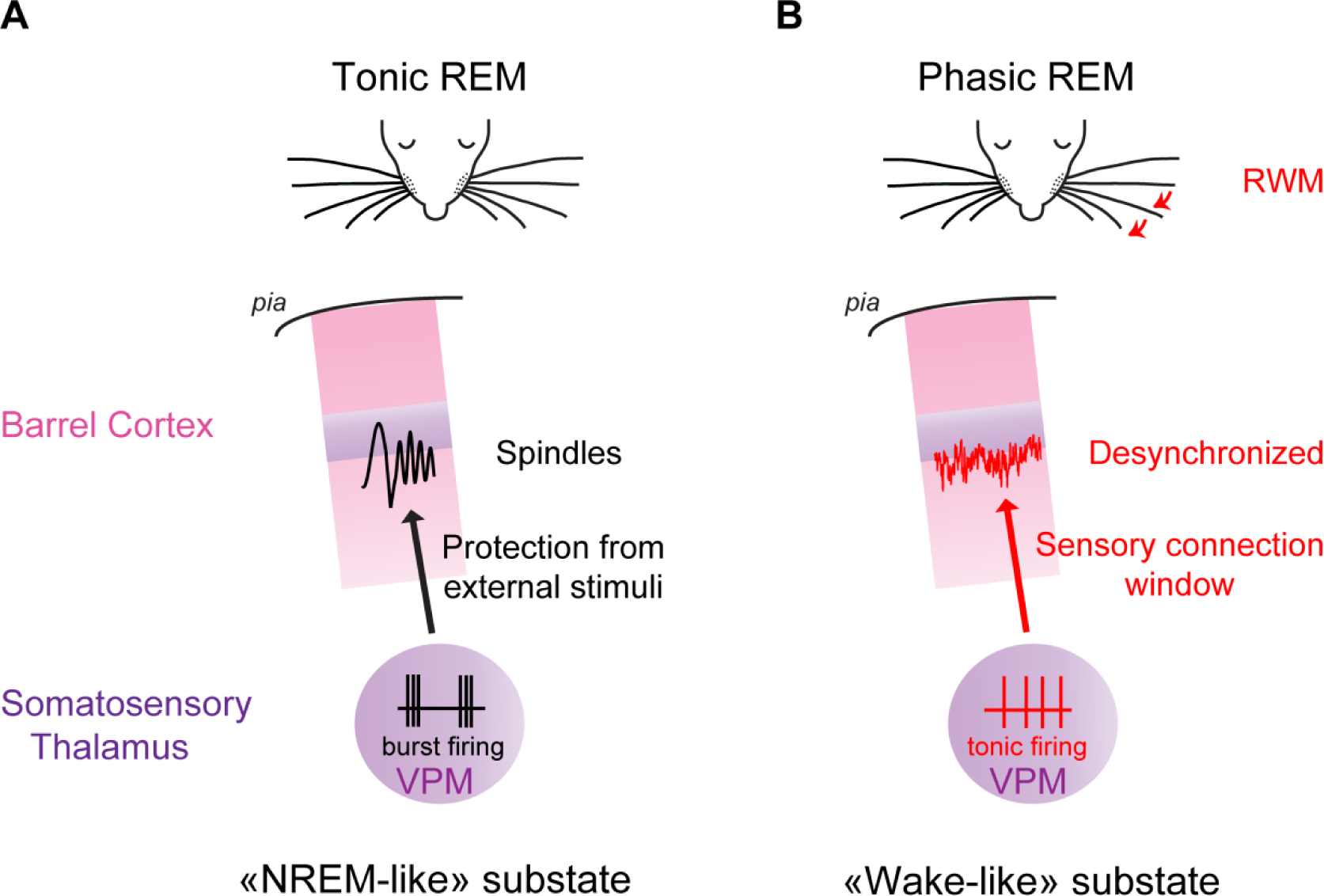
Differential thalamocortical dynamics during tonic and phasic states of REM. (A) During tonic REM, spindle bouts originating in the somatosensory thalamus spread to barrel cortex and thalamic neurons fire high frequency bursts, which may prevent efficient transfer of sensory information to the cortex. The system is in a NREM-like substate. (B) During phasic REM bouts, when whisker movements are observed, S1-LFP is desynchronized and thalamic neurons fire tonically, potentially allowing transmission of sensory information to the cortex. The system is in a wake-like substate.

## DISCUSSION

We report a transient cortical activation in S1-LFP associated with rapid whisker movements alternating with a high-amplitude and low frequency activity during REM, reminiscent of the S1-LFP activity observed in wakefulness and NREM respectively. By focusing on the lemniscal pathway of the well-defined mouse whisker system, we applied cross-correlations together with phase and time-frequency analysis to extra- and intracellular recordings performed in the thalamic VPM cells and the corresponding barrel somatosensory cortex and were able to demonstrate that some spindles activity propagating from VPM thalamus to barrel cortex is maintained during REM. We described rapid whisker movements during REM in adult mice which is of particular interest regarding REM sleep function considering that mice are nocturnal animals and extensively use their whiskers to explore their environment. RWM are faster and more foveal than those occurring during wakefulness (Figure 1), suggesting either that foveal whisking is more represented in REM or that RWM have different kinetics than wake whisking, as reported for eye movement properties in REM versus wakefulness ^46–49^.

During rapid whisker movements, we observed a cortical activation on S1-LFP similar to the one during whisking behavior in wakefulness; this activity alternated with a tonic state resembling NREM outside of these phasic events. However, both substates occurred during REM sleep episodes, clearly identified through neck muscle atonia and prominent EEG theta activity (Figure 1). RWM were associated with an increase in both theta frequency and gamma power measured on the EEG and HPC-LFP, as previously reported for rapid eye movements in humans and rodents ^10,36,50^. S1-LFP activity therefore appears highly dynamic across phasic and tonic REM substates. We also report here a strong delta power during the tonic REM, but restricted to S1 (Figure S2), in contrast to the high coherence of S1- and M1-LFP activity for delta oscillations reported in quiet wakefulness ^12,19^. Since functional coupling between distributed cortical areas through synchronized activities is thought to play an important role for many cognitive processes, including attention and decision-making, and support consciousness ^51–53^, especially in the delta-theta band ^54^, our findings suggest that cortical sensory networks process information in a more segregated manner during REM, potentially subtending the lack of consciousness during REM and dreaming.

We uncovered during REM, together with delta waves, the unexpected presence of spindle oscillations (Figure 2), which are typically associated with NREM. Given the thalamic origin of cortical NREM spindle activity ^40,45,55,56^, we looked for thalamic neuronal correlates of REM spindles observed on S1-LFP and found that a subset of VPM cells (16/37) maintained high frequency bursting activity while the animal was in REM; a firing mode typically associated with NREM. These VPM bursts of spikes displayed the same properties as those observed during NREM (Figure 3), including preferential firing on the rising phase of cortical spindle cycles ^40^. These results suggest that the thalamocortical mechanisms generating spindle activity during NREM are maintained during REM. However, we observed a depolarization of VPM-Vm in REM compared with Vm in NREM, as previously shown by Hirsch (1983) ^44^ in cats, the only paper in the literature to our knowledge reporting intracellular recordings in thalamic cells during REM. This finding raises the question: how can thalamic VPM cells promote spindle activity during REM if they are depolarized? It was previously shown that the activation of only a small fraction of the T-type channel population is required to generate robust calcium potentials in the ventrobasal thalamic nuclei *in vitro*, suggesting that high frequency bursts of action potentials can actually be evoked at depolarized potentials, when the vast majority of T-type channels are inactivated ^57^. Indeed, while Vm was on average was more depolarized during REM, we observed subthreshold oscillations in the spindle frequency range in a subset of VPM cells. Therefore, a more depolarized VPM-Vm would not prevent VPM cells from being enrolled by the thalamic reticular nucleus and, in turn, from entraining cortical spindle oscillations.

Our study indicates that, at the level of the barrel cortex, REM sleep is not a homogeneous sleep state, but rather consists of a mixture of NREM-like tonic periods interspersed with wake-like activity during phasic RWM. Tonic and phasic REM sleep may play different roles in information processing ^10,58^. Why the brain triggers RWM during sleep, and what it does with tactile feedbacks from these events remains a mystery. RWM, reminiscent in rodents of exploratory behavior in wakefulness, are in line with highly structured movements described in mammals during REM ^59,60^, or with the rapid changes in skin patterning and texture recently described in octopus ^61^. These behaviors all suggest a phasic expression during sleep of biologically and ethologically meaningful motor sequences, and may reflect globally orchestrated representation of virtual navigation ^59,62,63^.

Our data also reveal that S1-LFP and neuronal activity in its corresponding thalamic nucleus display unambiguous hallmarks of NREM during tonic REM, opening new avenue on the function of REM sleep. NREM-spindles were shown to elevate arousal threshold and protect the sleeper’s brain from external stimuli ^64^. While the same mechanism is likely to take place during REM spindle bouts, ongoing tonic firing activity in half of VPM neurons (Figure 3) may set the thalamocortical network at an “activatable state” level, rapidly allowing sensory transmission to resume during phasic episodes of RWM (Figure S4). Indeed, a recent study provided evidence in human of transient windows of sensory connection with the outside world during which external information can be processed in REM ^65^. Increased cholinergic activity in S1 may further act to suppress delta cortical activity during RWM ^66^, as reported during whisking in wakefulness ^67^. Alternatively, delta waves were described to support an activity-dependent process that can weaken reactivation of memory traces and promote forgetting ^68^. The inclusion of NREM features during REM may therefore subtend memory consolidation and forgetting of unnecessary material ^22–28^. The replay and transfer of recently acquired sensory information between neocortex and hippocampus is thought to take place during NREM, most especially during spindles, while memory is believed to be consolidated through hippocampal theta rhythms during REM sleep ^27,36^. The presence of spindle bouts during REM may reflect the opening of windows of communication between S1 and HPC and allow sensory neocortex to dynamically participate in hippocampal memory consolidation during REM sleep ^69^.

In summary, our findings show that at the level of sensory thalamus and barrel cortex, REM sleep is not a homogeneous sleep state, but rather consists of a mixture of NREM-like tonic periods interspersed with wake-like activity during phasic RWM. The NREM-like activity could potentially reduce transmission of bottom-up sensory information, contributing to brain disconnection from the outside world ^12,13^ and memory consolidation, while thalamocortical activation during RWM could facilitate the vivid internal sensory experiences that frequently occur during this state in the form of dreams ^70^.

## ACKNOWLEDGMENTS

We thank Sébastien Arthaud and the animal facility of the CRNL for technical support. We deeply thank Imola Mihalecz and C. Balazuc for technical assistance and help with the experiments. We warmly thank M. Bodier, N. Fourcaud-Trocmé, C. Loisel, A. Legay, O. Braud and A. Le Houssel for help with data processing and analysis. We are grateful to P. Salin and A. Hay for advices and critical reading of the manuscript. This work was funded by a ministerial doctoral grant (F.B.), ANR PARADOX (ANR-17-CE16-0024) (N.U. and L.G.), ANR Pom-Pom (ANR-21-CE16-0017) (N.U.).

## AUTHOR CONTRIBUTIONS

F.B. and N.U. performed the experiments. N.U. designed the project. F.B., K.J., L.G. and N.U. analyzed the data. T.D. helped with mice habituation and data processing. F.B. and. N.U. wrote the manuscript. L.G. commented on the manuscript.

## DECLARATION OF INTERESTS

The authors declare no competing interests.

## SUPPLEMENTAL INFORMATION

Supplemental information includes 4 figures and 3 tables.

## METHODS

### EXPERIMENTAL MODEL AND SUBJECT DETAILS

All procedures were approved by the “Ministère de la Recherche et de l’enseignement supérieur” (authorization APAFIS #35416) and were conducted in accordance with the European directive 86/609/EEC on the protection of animals used for experimental and scientific purposes. All efforts were made to minimize the number of animals used, as well as their stress and suffering. Experiments were carried out in male 8- to 16-week-old C57BL6J mice. After surgery, mice were housed individually to avoid deterioration of the implanted connector; they were weighted and handled daily. All animals were housed in standard conditions (23 ± 1 °C, food and water ad libitum) with toys to enrich their environment. All experiments were performed during the light part of the cycle (12 h light - 12 h dark).

### METHOD DETAILS

#### Surgery for head-fixation

Mice were anesthetized with a solution of ketamine (100 mg/kg) and xylazine (8 mg/kg) and were positioned in a stereotaxic frame. Body temperature was monitored and maintained at 37°C with an electric heating pad and eyes were covered with Liposic to prevent from ocular dryness. The skull was exposed, carefully cleaned with Betadine, and placed horizontally. Two stainless-steel screws were implanted over temporal and frontal cortical areas of the right hemisphere to measure the electroencephalogram (EEG, Figure 1A). Two flexible steel wires were inserted into neck muscles to monitor the electromyogram (EMG, Figure 1A). A light-weight metal head-holder, fixed to a micromanipulator fastened to the stereotaxic frame, was glued to the skull, as well as recordings chambers above the regions of interest (S1 and thalamus, M1 or hippocampus for a subset of mice) of the left hemisphere. The bone was then covered with a thin layer of acrylic cement (C&B Metabond). The metal piece was embedded in dental cement with the EEG screws, the EMG wires and their four-pin connector. To prevent infection, a thin layer of cyanoacrylate glue was applied on the exposed skull before sealing the recording chamber with silicon cement (Kwik Cast, WPI, USA). An iodine solution (Betadine 10%) was applied to all the borders of the implant for subsequent healing. Twenty minutes before the end of the surgery, the mouse received a subcutaneous injection of carprofen (5mg/kg). Animals were allowed to recover for five days before habituation sessions began. The implant (about 1g total weight, i.e., metal implant + connector + cement embedding) was well-tolerated by mice which were able to move, sleep, feed and drink normally in their home cage.

#### Mice training and habituation

During three to five weeks, repetitive daily training sessions of increasing durations were performed to habituate the mice to the head restraint. Their head was painlessly secured by screwing the metal piece, cemented to the mouse’s head to a head-fixation pillar; their body lying comfortably in a cardboard roll carpeted with soft tissue. At the end of the training period, mice remained calm for several hours, during which periods of quiet wakefulness, NREM and REM were typically observed.

After the training period and before the first recording session, a small craniotomy was performed, under isofluorane and buprenorphine (0.05 mg/kg), over the barrel cortex (S1; anteroposterior: −1.7mm relative to bregma, lateral: +3.5mm relative to midline; Paxinos and Franklin, 2001) in order to target LFP recordings from layer 5. Another craniotomy (about 1 mm) was drilled over the thalamus (anteroposterior: −1.7mm relative to bregma, lateral: +1.7mm relative to midline). For a subset of mice, a third craniotomy was performed over the primary motor cortex (M1; anteroposterior: +1mm relative to bregma, lateral: −1mm relative to midline) or the hippocampus (HPC; anteroposterior: −2mm relative to bregma, lateral: +1.2mm relative to midline). The dura mater was then removed in all craniotomies and the recording chambers were closed with Kwik Cast. All whiskers except for C2 were trimmed 10 to 15 mm from the skin. After three to four hours of recovery, the mouse was placed in the restraining frame as during the training sessions, and recordings were performed. In order to prevent eventual food or water deprivation during head-restraint, animals were fed regularly outside recording periods.

#### Single-unit and polygraphic recordings

LFP recordings (Grass QP511; band-pass filtered 0.1 - 300 Hz) were performed with glass micropipettes (6-8 µm tip diameter) filled with sodium chloride (0.15M) and lowered into layer 5 (−600 µm) of S1 barrel at 30° angle. In some experiments, glass micropipettes were vertically lowered in layer 6 (−1000 µm) of M1 and the CA1 region (−1200 µm) of the HPC.

Single-units in the thalamus were recorded using glass micropipettes (≤1 µm tip diameter, 30-80 MΩ) lowered vertically and filled with a solution of potassium acetate (0.5M) and Neurobiotin (2%, Vector Laboratories, Burlingame, CA, USA). The signal was amplified and low-pass filtered (10kHz, Cygnus, NeuroData), fed to an oscilloscope and an audio monitor, and digitized at 20 kHz. LFP and thalamic recordings were collected together with amplified EEG (band-pass filtered 0.1 - 300 Hz) and EMG (band-pass filtered 30 - 300 Hz) polygraphic signals (Grass QP511). The reference electrode for recordings with glass micropipettes was located on the surface of the skull, inside the recording chamber filled with sterile saline solution. All signals were collected via a CED power 1401 analog-to-digital converter (Cambridge electronic design, UK) using Spike2 software.

Initial localization of thalamic cells was based on the coordinates described in Paxinos and Franklin (2001). Manual deflection of individual contralateral (right side) whiskers was performed in order to determine the receptive field of recorded cells. All cells assigned to the VPM and tested for whisker deflections were mono- or multi-whisker responsive cells. Cell locations were determined by the juxta- or intra-cellular labelling of the last recorded cell (Pinaut, 1996) and the reconstruction of each pipette track.

At the end of experiments, mice were perfused under deep anesthesia with saline, followed by a fixative containing 4% paraformaldehyde in phosphate buffer (PB 0.1M, pH=7.3). Brains were post-fixed for 1h, and coronal slices were subsequently cut at 50 µm thickness with a vibratome. Sections were processed for cytochrome oxidase and Neurobiotin histochemistry according to standard protocols (Veinante et al., 2000).

### QUANTIFICATION AND STATISTICAL ANALYSIS

#### Cortical and behavioral states

The activity of thalamic cells was recorded together with EEG and LFP in the ipsilateral S1 (C2 column) in layer 5 (600mm depth). In addition, the muscle tone was monitored from the neck (EMG) and whisker movements were tracked with a high-speed camera. All these parameters allowed us to determine the vigilance state of the mouse. In this paper, we focus on cortical and thalamic activity during REM and its relationship with other vigilance states of sleep and wakefulness, specifically non-REM sleep (NREM), Whisking (Wh), active and quiet wakefulness (respectively aWK and qWK; Urbain et al., 2015, 2019 ^18,40^). REM was characterized by the absence of EMG tone and a prominent peak in the theta range (6 - 9 Hz) in the EEG. Only the presence of whisker movements, or not, distinguished RWM from REM-noWh. In REM, whisking episodes were considered when whisker movements were at least 15° in amplitude and lasted at least 300ms. Periods of REM with either jaw chattering, low amplitude (inferior to 15°) or short duration whisker movements (less than 300ms) were not scored neither as RWM nor REM-noWh (about one third of REM). NREM was distinguished by high-amplitude, low frequency EEG and LFP (0.1 - 1.5 Hz) activity with a strong dominant frequency range of 10 - 18 Hz (spindle activity, see Figures 2E), associated with a weak EMG tone. NREM is clearly distinguishable from REM, since REM is characterized by cortical EEG spectra with a strong dominant theta band (6 - 9 Hz) and no muscle tone at EMG. Quiet wakefulness was distinguished by high-amplitude, low frequency (2 - 8 Hz) EEG and LFP activity associated with a weak EMG tone. Importantly, NREM is a different state from qWK: both cortical LFP and EEG spectra are clearly distinct in NREM and qWK, with a strong dominant frequency range of 2 - 6 Hz in the latter state (see Figure S2H; Urbain et al., 2015, 2019 ^18,40^). Active wakefulness and Whisking were characterized by lower amplitude and higher frequency EEG and LFP oscillations than quiet wakefulness, associated with sustained EMG activity; polygraphic features of the Active and Whisking states are similar, but in the latter case the mouse was whisking. Whisking episodes in wakefulness were considered when whisker movements were at least 20° in amplitude and lasted at least 300ms; periods of time associated with whisker movements of smaller amplitude or shorter duration were excluded from the analysis. Transitional and intermediate states (such as drowsiness or transition from NREM to REM sleep) were excluded from the analysis. Spike time occurrences were determined as threshold amplitude crossings. The spike detection was checked (and corrected if necessary) for each episode included in the analyses. An automatic scoring of these states was performed using a custom-written software in Matlab and the detection was systematically checked manually.

#### Quantification of whisker movements

Electrophysiological recordings were performed while the whiskers movements of the mouse were simultaneously filmed using a high-speed camera (Photon Focus MV1-D2048x1088I-96-G2) operating at 400 frames per second. Continuous recordings were performed in 600 seconds bouts. Whisker movements were quantified off-line using DeepLabCut interface and custom written routines running with Python to track whisker position, measured as the angle between the nose, the attachment of the whisker to the whisker pad and a point on the whisker. Whisking occurrences were found using a custom written routine running with MatLab and checked manually. Whisking amplitude of each episode was defined as the difference between the maximum whisker position and the mean angle 200 ms before. To optimize the detection of whisking cycles, whisker position was high-pass filtered at 2Hz. Whisking cycles were then detected using the function “*findpeaks*” of Matlab, which also returned the prominence of each peak (see figure 1E). Note that a minimum peak prominence ‘MinPeakProminence’ was set at 5° to optimize the detection in both wakefulness and REM. However, due to the whiking kinematic in REM, described in the Results part and illustrated in the Figure 1C, we were not able to detect all whisking cycles in REM. The number of cycles / whisking episod is therefore underestimated and the mean cycle prominence overestimated in REM compared with those measured in wakefulness.

#### Spectral analysis

Raw data were imported from spike2 to MATLAB and processed using custom-written functions. EEG and cortical LFPs were filtered (1 - 256 Hz) and down-sampled to 512 Hz. Signals were classified per vigilance state and 1-second epochs were selected in each state. All epochs were then collected for each mouse and each state. A Hanning window was applied to each epoch and all the analysis described in this section were computed on these 1s epochs, precluding us from studying oscillations below 2 Hz, but allowing us to analyze short phasic events such as whisking and RWM. All epochs were averaged to get the result per mouse and per state. Power spectral density (PSD) was computed using the *mtspectrumc* function of the chronux toolbox and the spectral power in the different frequency ranges was calculated as the sum of the PSD in the range of interest. Normalized PSD (Figure S2) were computed by dividing the PSD by its sum between 1 and 45 Hz. Theta peak frequency was computed, for each mouse, as the frequency of the maximum value of the PSD between 5 and 12 Hz. From this value *fmax*, theta peak spectral power was computed in the range [*fmax*-2Hz : *fmax*+2Hz]. The correlation between the EEG and the HPC-LFP was computed using the functions *xcorr* in Matlab. In addition, we computed the correlation between the HPC- LFP and the randomly permuted EEG using the function *circshift (surrogates).* This step was repeated 100 times. To assess the statistical significance of the average correlation coefficient for each mouse, we compared the amplitude of the real correlation coefficients with those of the surrogates.

In order to analyze theta/gamma modulation, theta and gamma oscillations were extracted from each signal using the Ensemble Empirical Mode Decomposition (EEMD). This method decomposes the signal into so-called intrinsic mode functions (IMF) that correspond to a time series of a frequency present in the signal. Selecting the IMFs with a maximum frequency in the theta (5 - 12 Hz) and gamma (60 - 90 Hz) ranges allowed us extract theta and gamma oscillations without any modifications of the raw data (no filter). Theta gamma comodulograms (Figure S1F) were examined by computing the coherence between the signal (EEG or HPC-LFP) and gamma power time series for each frequency (bin 1Hz; see Sirota et al., 2008 ^39^). Phase analysis (Figure S1G) was performed following the method described by Tort et al. (2010): first, the time series of the theta oscillation phases were obtained using the Hilbert transform and the envelop of the gamma oscillation was extracted. Then, theta phase was binned (5° bins) and the mean amplitude of the gamma composite envelope was calculated on each bin. Finally, the mean amplitude was normalized by the sum of the amplitude across all bins. The histogram obtained with this method was fitted with a sine wave for each mouse in order to compute the modulation index (max-min of the sinusoidal fit) and the preferred phase of gamma oscillation (phase of the sine wave peak; see polar plots). Gamma envelops were also randomly permuted using the function *circshift* in Matlab (surrogates) to compute the mean amplitude of gamma surrogates on each bin of theta phase. To assess the statistical significance of theta/gamma modulation, a Wilcoxon test was performed between the modulation index of the raw data and the surrogates.

#### Spiking analysis

Spike time occurrences were determined as threshold amplitude crossings on Spike 2 and spike detection was checked for all recordings included in the analysis. Mean firing rate was computed per episode for each state, then averaged across all episodes for each cell. To assess whether a cell was significantly modulated by whisking behavior in wakefulness and/or in REM, we selected periods of active wakefulness or REM- noWh occurring 1s before or after the whisking episode, and we then compared its firing rate across these paired episodes of Wh/aWK and RWM/REM-noWh. Only cells for which at least five paired episodes were recorded were included in this analysis (t-test for paired data, significant if p<0.05).

Bursts were defined as two or more spikes that were preceded by at least 65ms of silence and had interspike intervals (ISIs) < 8 ms. A maximal ISI of 8 ms was chosen in order to include the last spikes fired in long bursts ^40^ (Urbain et al., 2019; see Figure S3). Only cells for which > 100 spikes were collected in the states of interest (NREM, REM-noWh and RWM for the study of the percentage of spikes included in bursts, NREM and REM-noWh for the comparison of bursts characteristics) were considered for this study. Cells were considered to fire in bursts in one state if the percentage of spikes included in bursts was superior to 50%. No ISI < 1ms was observed (Figure S3), indicating single-unit recording. In order to illustrate the high frequency bursts activity during NREM and REM, we showed on Figure 3D the instantaneous frequency of the thalamic cell, calculated as 1/interspike interval (ISI).

#### Spindles detection

S1-LFP spindles were observed as transient waxing and waning oscillatory events in the 10 - 18 Hz range (see Methods in Urbain et al., 2019 ^40^). Briefly, S1-LFP spindles were detected by applying a ridge line detection on the wavelet time frequency map using a custom-written script in Python. For spindles detection during NREM, the threshold was set as 3 times the median power of the 10 - 18 Hz band excluding NREM periods. This threshold was adjusted from our previous study (threshold of 4) in order to take into account the decrease in REM spindles amplitude compared to NREM (Figure S2F). Only oscillations spanning more than 3 cycles were kept for analysis. Each detected spindle was converted into a time frequency line giving for each time point: instantaneous frequency, phase and power of the spindle. For comparison of spindles parameters between NREM and REM, only spindles in NREM episodes preceding REM were considered (in order to have the same number of NREM and REM episodes in both samples) and a minimum of 5 spindles in each state was required to be considered for this analysis.

#### Phase analysis

Phase analysis of VPM neurons firing onto S1-LFP spindles was performed following the method already used in our previous paper (Urbain et al., 2019 ^40^). The phase of action potentials (APs) relative to spindles in the S1-LFP was determined by the angle of the instantaneous phase (derived from a Hilbert transformation of the S1-LFP) at the location of spikes. The trough of the spindles was defined as 0° and the peak as 180°. These two points define the reference frame, which enabled us to compare waves with different shapes and lengths. For each cell, we built phase histograms of spike timing relative to spindle troughs. Only neurons which fired at least 50 action potentials during spindle bouts were included in the analysis. Firing was considered to be modulated by the oscillations if p< 0.01 using the Rayleigh’s test. To compute the grand average of modulated cells (see Figure 4), the number of spikes per phase bin was normalized for each cell (by dividing by the total number of spikes over one spindle cycle).

Thalamic spikes were collected and aligned at their peaks to compute S1-LFP spike-triggered averages (STA). Only spikes separated by more than 65 ms were considered. Only cells with at least 50 spikes in both NREM and REM were kept for the analysis. S1-LFPs were collected and averaged per cell (436.7 ± 51.3 spikes per cell in NREM, 236.3 ± 28.3 spikes per cell in REM; range: 83 - 1333 spikes per cell in NREM, 61 - 734 spikes per cell in REM), then a grand average was calculated with the sem. In addition, we computed the STA of the randomly permuted S1-LFP using the function *cricshift* of Matlab (surrogates). For each VPM spike, S1-LFP was permuted 100 times in the interval −0.1 to 0.2s relative to the spike. The average surrogate STA for the cell was the average of all the permutations on all spikes. To assess the statistical significance of the STA in each state (NREM and REM), the peak value of S1-LFP and its surrogates was computed for each cell and a Wilcoxon test was performed.

For the VPM spikes distribution on the delta waves, spindles preceded by a delta event were detected by visual inspection for each cell, and aligned at the spindle onset. Thalamic spike time occurrences were distributed in a Peristimulus Time Histogram (−500 to 500 ms PSTH, 50 ms bins, see Figure 4F). Only the time windows for which the cell fired at least 10 spikes were kept for the analysis, and only cells with at least 5 windows (i.e., 5 delta events with at least 10 spikes) were considered.

#### Statistical analysis

Data analysis was performed using Spike2, MATLAB, Python and Excel software. Unless otherwise stated, results are reported as mean ± SEM. Statistical analysis was performed using Matlab. The Tukey-Kramer or the Dunn-Sidak test was chosen when a post hoc test was applied. Circular statistics was performed using the “*circular statistics toolbox*” in Matlab. Statistical tests are specified in the figure legends. The numbers of cells and mice considered for each test are provided in the main text or / and in figure legends. Additional statistical measures (such as mean, median, range) are provided in Tables S1, S2 and S3. Except otherwise stated, spectral power for EEG and LFP data were compared across states using Friedman or Wilcoxon tests; other data were compared across states using ANOVA for repeated measures or *t*-test for paired data. Significance levels *p< 0.05; **p< 0.01; ***p< 0.001.

---

**Table S1:** EEG (no color) and hippocampal LFP (HPC-LFP, green) activity in the theta and gamma bands (mean ± SEM and median with range) calculated from EEG and HPC-LFP recordings in REM (without, REM-NoWH or with whisking, RWM). Statistical comparisons available in Figures 1 and S1.

|  | REM-noWh | RWM | Nb of mice |
| --- | --- | --- | --- |
| EEG theta peak frequency (Hz) | 6.7 ± 0.1 | 8.8 ± 0.1 | 32 mice recorded across all states |
| Median (min - max) | 7 (6 - 8) | 9 (7 - 10) |  |
| EEG theta power (μV<sup>2</sup>) | 1.2 ± 0.1 | 1.2 ± 0.1 | 32 |
| Median (min - max) | 1.2 (0.2 - 3.2) | 1.2 (0.2 - 2.8) |  |
| HPC-LFP theta peak frequency (Hz) | 7 | 9 | 3 mice with both HPC-LFP and EEG recordings |
| Median (min - max) | 7 (7 - 7) | 9 (9 - 9) |  |
| HPC-LFP theta power (μV<sup>2</sup>) | 3.4 ± 0.2 | 3.7 ± 0.2 | 3 |
| Median (min - max) | 3.6 (3.0 - 3.7) | 3.8 (3.2 - 3.9) |  |
| EEG / HPC-LFP correlation | -0.66 ± 0.06 | -0.71 ± 0.04 | 3 |
| Median (min - max) | -0.64 (-0.77 to 0.57) | -0.71 (-0.78 to -0.65) |  |
| EEG gamma (60 – 90 Hz) power (μV<sup>2</sup>) | 0.036 ± 0.003 | 0.056 ± 0.004 | 23 out of 32 mice analyzed for gamma |
| Median (min - max) | 0.034 (0.008 - 0.065) | 0.058 (0.024 - 0.089) |  |
| HPC-LFP gamma (60 – 90 Hz) power (μV<sup>2</sup>) | 0.24 ± 0.04 | 0.44 ± 0.11 | 3 |
| Median (min - max) | 0.27 (0.16 - 0.30) | 0.50 (0.22 - 0.59) |  |
| EEG Theta phase for maximum gamma amplitude (°) (0°: Theta trough) | 6.7 ± 5.5 | -0.5 ± 6.0 | 23 |
| Median (min - max) | 5 (0 - 355) | 0 (0 - 355) |  |

**Table S2:**
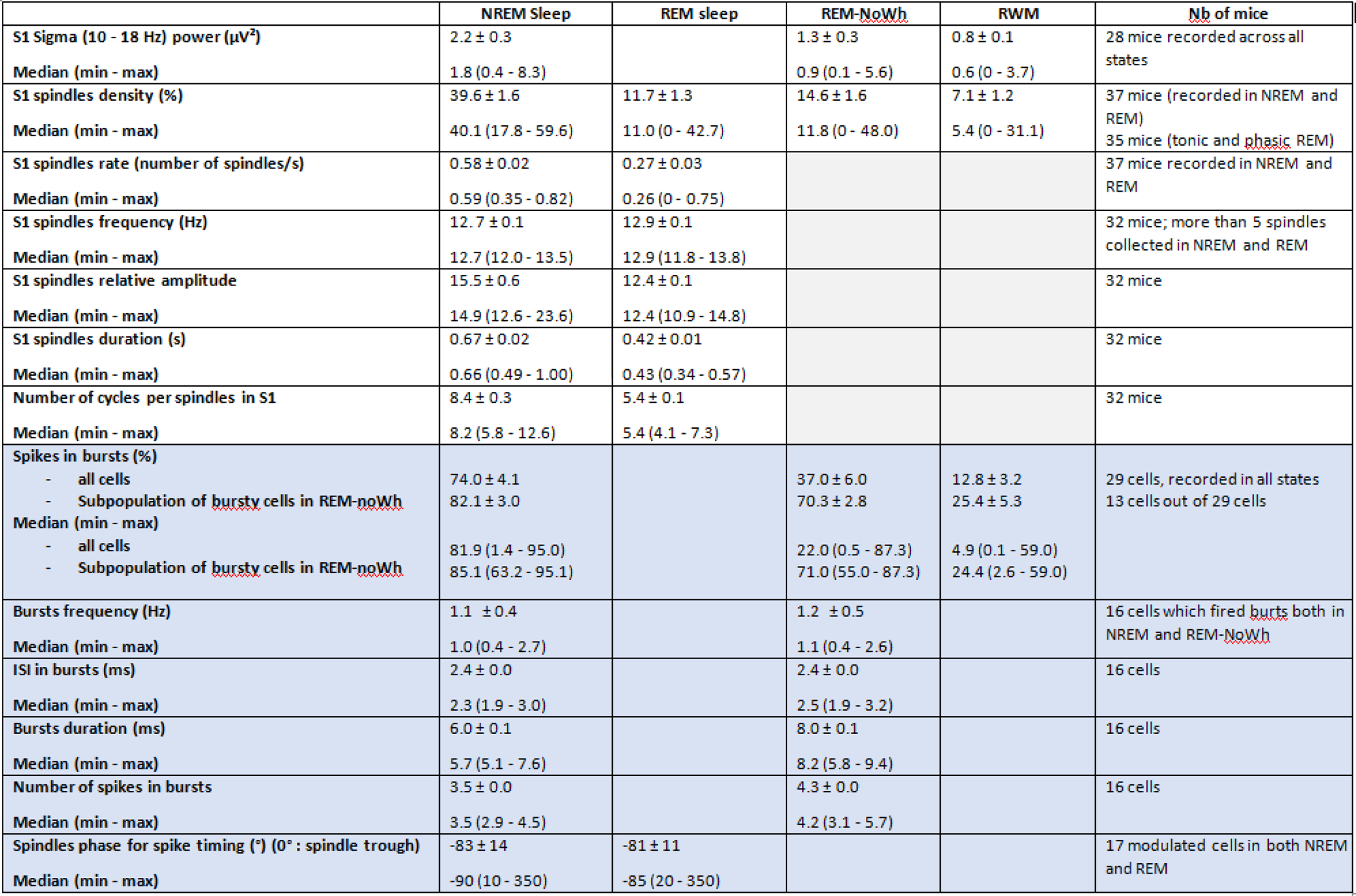
Spindle features (no color) and thalamic firing (blue) (mean ± SEM and median with range) calculated from S1-LFP and thalamic extracellular recordings in NREM and REM sleep. Statistical comparisons available in Figures 2, 3, and S2.

**Table S3.**
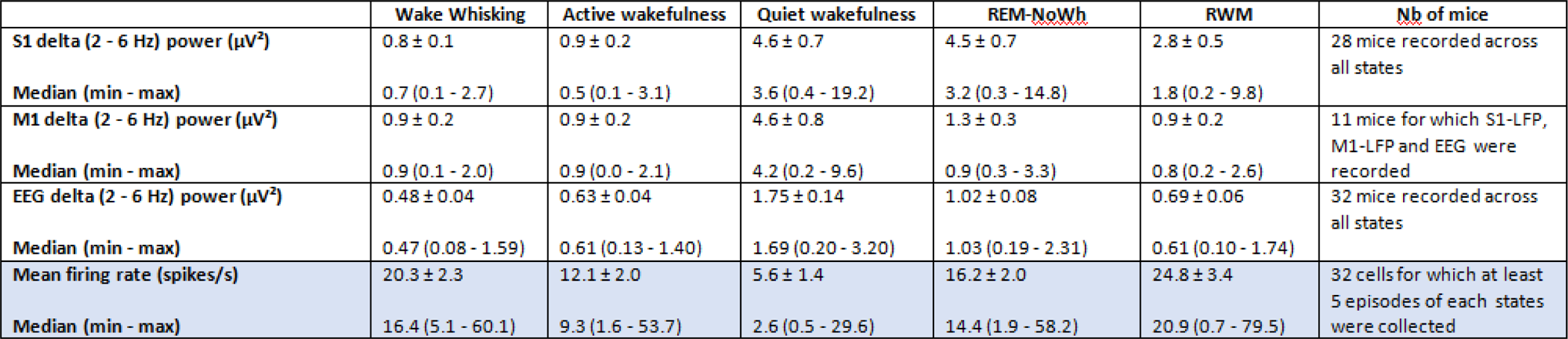
Cortical delta power (no color) and thalamic firing rate (blue) (mean ± SEM and median with range) calculated from S1-LFP, M1-LFP and EEG, and thalamic units recordings across the sleep-wake cycle (REM-NoWH, REM without whisking or tonic REM; RWM, REM with whisking or phasic REM). Statistical comparisons available in Figures 2, 3 and S2.

|  | Wake Whisking | Active wakefulness | Quiet wakefulness | REM-NoWh | RWM | Nb of mice |
| --- | --- | --- | --- | --- | --- | --- |
| S1 delta (2 - 6 Hz) power ($\mu V^2$) | 0.8 ± 0.1 | 0.9 ± 0.2 | 4.6 ± 0.7 | 4.5 ± 0.7 | 2.8 ± 0.5 | 28 mice recorded across all states |
| Median (min - max) | 0.7 (0.1 - 2.7) | 0.5 (0.1 - 3.1) | 3.6 (0.4 - 19.2) | 3.2 (0.3 - 14.8) | 1.8 (0.2 - 9.8) |  |
| M1 delta (2 - 6 Hz) power ($\mu V^2$) | 0.9 ± 0.2 | 0.9 ± 0.2 | 4.6 ± 0.8 | 1.3 ± 0.3 | 0.9 ± 0.2 | 11 mice for which S1-LFP, M1-LFP and EEG were recorded |
| Median (min - max) | 0.9 (0.1 - 2.0) | 0.9 (0.0 - 2.1) | 4.2 (0.2 - 9.6) | 0.9 (0.3 - 3.3) | 0.8 (0.2 - 2.6) |  |
| EEG delta (2 - 6 Hz) power ($\mu V^2$) | 0.48 ± 0.04 | 0.63 ± 0.04 | 1.75 ± 0.14 | 1.02 ± 0.08 | 0.69 ± 0.06 | 32 mice recorded across all states |
| Median (min - max) | 0.47 (0.08 - 1.59) | 0.61 (0.13 - 1.40) | 1.69 (0.20 - 3.20) | 1.03 (0.19 - 2.31) | 0.61 (0.10 - 1.74) |  |
| Mean firing rate (spikes/s) | 20.3 ± 2.3 | 12.1 ± 2.0 | 5.6 ± 1.4 | 16.2 ± 2.0 | 24.8 ± 3.4 | 32 cells for which at least 5 episodes of each states were collected |
| Median (min - max) | 16.4 (5.1 - 60.1) | 9.3 (1.6 - 53.7) | 2.6 (0.5 - 29.6) | 14.4 (1.9 - 58.2) | 20.9 (0.7 - 79.5) |  |

## Notes

### Competing Interest Statement

The authors have declared no competing interest.

